# C57BL/6 BAC-CAG Huntington’s disease mice show somatic CAG expansion and responses to small interfering RNAs comparable to the FVB strain

**DOI:** 10.64898/2026.05.08.723329

**Authors:** Jillian Belgrad, Ashley Summers, Samuel Hildebrand, Ellen Sapp, Eric Luu, Nozomi Yamada, Dan O’Reilly, Thomas F. Vogt, David Howland, X. William Yang, Marian DiFiglia, Neil Aronin, Anastasia Khvorova

## Abstract

Huntington’s disease (HD) is a neurodegenerative disorder caused by CAG repeat expansion in the huntingtin (HTT) gene, with longer repeats linked to earlier onset. Somatic CAG expansion, particularly in the striatum, contributes to disease progression and is influenced by HTT biology and genetic modifiers. Modulating somatic expansion is emerging as a promising approach to slow or prevent HD, and mouse models have been crucial for preclinical testing of different therapeutic strategies. The BAC-CAG model, developed on the FVB strain, has been used to study somatic expansion of human expanded HTT. However, comparisons with other key HD mouse models have been limited by differences in genetic background, as many other models are on the C57BL/6 strain. The BAC-CAG model has now been developed on a C57BL/6 background. To determine whether the C57BL/6 BAC-CAG model can be used to study and modulate somatic expansion, we compared CAG expansion in mice on C57BL/6 or FVB backgrounds, with and without intraventricular divalent small interfering RNAs (siRNA) targeting HD modifiers MutS homolog 3 (MSH3) and HTT. Both strains exhibited robust, comparable somatic expansion over two months, which was blocked by MSH3-, but not HTT-, targeted siRNA. RNA sequencing identified gene expression differences primarily in pseudogenes, with no differences in endogenous *Htt*, human *HTT*, or mismatch repair genes. These results demonstrate that BAC-CAG mice on a C57BL/6 background exhibit somatic CAG expansion comparable to the validated FVB strain, providing a model to study and preclinically test therapies targeting somatic expansion in HD.

## INTRODUCTION

Huntington’s disease (HD) is a progressive neurodegenerative disorder caused by CAG repeat expansion in the HTT gene, with longer repeats associated with earlier disease onset.^1-6^ Somatic CAG expansion, particularly in the striatum, drives disease progression and is influenced by HTT biology and genetic modifiers.^2, 3, 5, 7^ Genome-wide association studies have identified several such modifiers, most notably DNA mismatch repair genes such as MutS homolog 3 (MSH3), that alter the age of onset beyond that of the inherited repeat length.^3-5, 8^ Preclinical mouse studies support a central role for somatic expansion in HD pathogenesis, contributing to nuclear dysfunction, mRNA and protein aggregation, early neuronal impairment, and cell death.^3, 4, 6, 8-26^ Modeling somatic expansion in mouse models is therefore critical in understanding mechanisms of HD progression and developing therapeutic strategies to potentially halt or prevent HD pathogenesis.

Several HD mouse models exhibit robust somatic expansion, including Q111, Q140, Q175, and BAC-CAG (∼120 CAGs) lines, many of which are on the C57BL6 background strain, except for BAC-CAG.^9, 10, 14, 20, 21, 27-35^ BAC-CAG is a transgenic model carrying a bacterial artificial chromosome (BAC) with the human expanded *HTT* gene on an FVB background.^36^ The BAC-CAG model features the full human *HTT* gene, under human regulatory elements, with an uninterrupted expanded CAG region (120 CAGs) including the repeat structure (120 CAGs-1 CAA-8 CAGs-CAA-CAG) ^36^

BAC-CAG models recapitulate key HD features, observed across other HD mouse models, including robust somatic expansion, transcriptional dysregulation, protein aggregation, and behavioral deficits.^18, 27-30, 33, 35-37^ However, direct comparisons of biology and phenotypes between these BAC-CAG and other HD models are complicated by the known strong influence of genetic background on somatic expansion, particularly in the striatum.^10, 31^ While these models share many HD features, it also remains unclear which phenotypes reflect differences between the mouse *Htt* and human *HTT* gene structure versus the mouse genetic strain background. To address this confounding factor, the BAC-CAG model has now been developed on a C57BL/6 background. The goal of this study was to characterize the utility of the BAC-CAG C57BL/6 stain as a model to the study of somatic expansion and therapeutic testing in HD compared to the existing FVB model strain.

Therapeutic strategies for HD include modulation of somatic CAG repeat expansion, commonly through targeting MSH3, as well as reduction of HTT expression, which has been demonstrated in multiple model systems.^7^ siRNAs are an emerging class of nucleic acid-based therapeutics that modulate gene expression by selectively cleaving target mRNA.^38-40^ Fully chemically stabilized divalent siRNA enable durable, effective silencing of genes throughout the central nervous system (CNS) following intraventricular administration.^41^

To address the confounding effects of genetic background on somatic expansion, BAC-CAG mice were generated on a C57BL/6 background, aligning with other HD mouse models, and compared with the BAC-CAG model on the FVB background. We measured striatal CAG expansion in both strains, with and without siRNA targeting the HD modifiers *Msh3* and *HTT*. Both models showed robust somatic expansion over two months. In both strains, expansion was blocked by siRNA targeting *Msh3*, while *HTT* lowering had no effect. RNA sequencing showed no differences in endogenous *Htt*, mutant *HTT* or the DNA mismatch repair pathway between strains. These results demonstrate that BAC-CAG mice on the C57BL/6 background undergo somatic expansion as well as the validated FVB strain and offers a robust model for testing and developing therapies targeting somatic CAG expansion in HD.

## METHODS AND METHODS

### Solid-Phase Synthesis of Oligonucleotides

Oligonucleotides were synthesized via standard phosphoramidite chemistry on a MerMade12 automated synthesizer (Biosearch Technologies, Novato, CA). Phosphoramidites used included 2′-O-methyl (2′OMe) and 2′-fluoro (2′F) nucleotides with conventional protecting groups. The 5′-(E)-Vinyl tetraphosphonate (pivaloyloxymethyl) 2′-O-methyl-uridine 3′-CE phosphoramidite was incorporated for 5′-(E)-Vinyl phosphonate addition. Phosphoramidites were prepared at 0.10 M in anhydrous acetonitrile (ACN), except 2′-OMe-uridine, which was dissolved in ACN containing 15% dimethylformamide. Phosphoramidites were obtained from Hongene Biotech (Union City, CA) and Chemgenes (Wilmington, MA), and other reagents from Chemgenes. Solid supports consisted of LCAA-controlled pore glass with a 500Å Unylinker terminus (Chemgenes) or custom 500Å di-trityl protected supports linked via tetraethylene glycol (Hongene).

### Oligonucleotide Cleavage, Deprotection, and Purification

Oligonucleotides were cleaved from solid supports and deprotected. 5′-(E)-Vinyl phosphonate-containing oligonucleotides were treated with 3% diethylamine in ammonium hydroxide at 35°C for 20 hours under gentle agitation. Divalent oligonucleotides were deprotected using a 1:1 mixture of 40% aqueous monomethylamine and ammonium hydroxide at 25°C for 2 hours. After filtration and washing with 5% ACN in water, oligonucleotides were dried via centrifugal vacuum. Purification was conducted on an Agilent 1290 Infinity II HPLC system using Source 15Q anion exchange resin with a 30-70% linear gradient over 30 minutes at 50°C, monitoring absorbance at 260 nm. Selected fractions were pooled and desalted using Sephadex G-25 size-exclusion chromatography on an Akta FPLC. Purity and identity were confirmed by IP-RP analysis on an Agilent 6530 Accurate-Mass Q-TOF.

### Divalent siRNA Duplexing and Dosing

In vivo-grade siRNA was duplexed at 10 nmol per 10 µL (2000 nM). To mitigate acute neurotoxicity, siRNAs were prepared in calcium- and magnesium-supplemented artificial cerebrospinal fluid (aCSF).^42^ The sequences and chemical modification patterns of non-targeting control (NTC), MSH3 and HTT siRNA can be found in Supplemental Table 1.

### Mouse Work

All procedures were approved by the Institutional Animal Care and Use Committee at UMass Chan Medical School (IACUC protocol 202000010). Mice were bred and delivered from Jackson Labs (Farmington, CT) and transferred at 6 weeks to pathogen-free facilities. Animals were housed under a 12-hour light/dark cycle at 23 ± 1°C and 50 ± 20% humidity, with free access to food and water. Genotypes used included BAC-CAG on the FVB line (CHDI Stock 412730) and BAC-CAG on the C57B6 background (CHDI Stock 41608).

### Stereotaxic Intracerebroventricular Injections

Mice were anesthetized with isoflurane, and the scalp was shaved and incised. Burr holes were drilled using coordinates relative to bregma: mediolateral ±1 mm, posterior -0.2 mm, ventral -2.5 mm. Five microliters per ventricle were delivered at 750 nL/min bilaterally. Post-injection, mice received Meloxicam ER and saline and were monitored daily for 72 hours and weekly thereafter. Mice were injected at age three months and sacrificed at age five months, two months post injection.

### mRNA Quantification

At sacrifice, 1.5 mm × 1 mm brain punches were collected across regions and stored in RNAlater overnight. Punches were lysed in 600 µL homogenizing buffer (Invitrogen QG0517), and mRNA was quantified using the branched DNA Quantigene Singleplex assay (Invitrogen QS0016) using the following probes: *Fan1* (Mouse, SA-3051642), *Htt* (Mouse, SB-14150), *Mlh1* (Mouse, SA-3030156), *Mlh3* (Mouse, SB-3046475), *Msh2* (Mouse, SA-3030207), *Msh3* (Mouse, SA-3030208), *Msh6* (Mouse, SA-3030210), *Pms1* (SA-3046971), *PMS2* (Human, SA-3002748), and *Pms2* (Mouse, SA-3030786) and Hprt (Mouse, SB-15463) for housekeeping controls. Expression levels were normalized to housekeeping genes and treatment efficacy quantified relative to non-targeting controls.

### Protein Preparation and Western Blotting

Frozen tissue punches were homogenized on ice in 10 mM HEPES pH 7.2, 250 mM sucrose, 1 mM EDTA, protease inhibitors, 1 mM NaF, and 1 mM Na3VO4, followed by 10 s sonication. Protein concentrations were measured via Bradford assay (BioRad). 10–20 µg protein per sample were separated on 3–8% Tris-acetate SDS-PAGE gels (BioRad) and transferred to nitrocellulose (TransBlot Turbo, BioRad). Blots were probed with antibodies against HTT (aa1–17, 1:2000), MSH3 (1:500, Santa Cruz sc-271079), tubulin (1:4000, Sigma T8328). Bands were visualized with ChemiDoc XRS+, and area and intensity quantified in ImageJ. Total signals were normalized to tubulin. Statistical outliers were removed with ROUT analysis. Statistical significance was assessed via one-way ANOVA with Tukey’s post hoc tests.

### Fragment Analysis PCR

1.5 mm × 1 mm striatal or medial cortex punches were flash-frozen. DNA was extracted using Qiagen DNeasy Blood & Tissue Kit (Cat. 69506). Fragment analysis PCR was performed with AmpliTaq 360 DNA Polymerase (ThermoFisher Cat. 4398818) using 150 ng DNA per reaction. Primers: forward CAG1 6FAM-ATG AAG GCC TTC GAG TCC CTC AAG TCC TTC; reverse HU3 GGC GGC TGA GGA AGC TGA GGA. Thermocycling: 94°C 90 s; [94°C 30 s; 63.4°C 30 s; 72°C 90 s] × 35; 72°C 10 min. Fragment sizes were determined with GeneScan LIZ 500 ladder and analyzed with ThermoFisher PeakScanner software. Somatic instability was quantified using the Lee et al. instability index.^43^

### RNA Sequencing and Transcriptomic Analysis

At sacrifice, 1.5 mm × 1 mm striatal punches were flash-frozen. RNA was extracted from one punch per NTC-treated mouse using the Monarch Total RNA Miniprep Kit (T2010S), and quality was assessed with the Agilent RNA ScreenTape (#5067). Sample size was N=5 mice per strain. Purified RNA samples were submitted to Genewiz for bulk RNA sequencing. Sequencing reads were aligned to the mouse reference genome using Kallisto (mm10). Normalized counts and adjusted p-values were calculated using DESeq2.

### Statistics

All statistics were performed in GraphPad Prism v10.2.3. One-way ANOVA with Tukey’s post hoc test was used for single-variable comparisons of more than two groups, and two-way ANOVA with multiple comparisons for experiments with two variables. Outliers were identified and removed with ROUT analysis. RNA-sequencing data was analyzed using DESeq2-adjusted p-values.

## RESULTS

HD is characterized by mutant HTT expression and progressive neuronal degeneration, features that must be modeled to support therapeutic development. The BAC-CAG model, first developed by the Yang lab on the FVB strain, contains the full-length human HTT sequence with 120 CAGs and recapitulates several core aspects of disease biology including protein aggregation, neurodegeneration, sleep deficits.^36^

To assess the impact of background strain on BAC-CAG somatic expansion and response to therapeutic modulation of HD modified, we probed the response to divalent siRNAs targeting *Msh3* or *HTT* in BAC-CAG mice on FVB and C57BL/6 backgrounds. Mice were injected intracerebroventricularly at three months of age with non-targeting control (NTC), *Msh3*-targeting, or *HTT*-targeting siRNA. At five months of age, mRNA levels, protein levels, and somatic CAG repeat expansion were evaluated.

Two-months post injection, MSH3-targeted siRNA reduced *Msh3* mRNA to ∼20-80% of NTC in FVB mice and 20-40% in C57BL/6 mice in the striatum and medial cortex (Figure 1A, two-way ANOVA vs NTC, p < 0.001). HTT-siRNA reduced mutant *HTT* mRNA by ∼50-70% in both strains (Figure 1B, p < 0.0001). mRNA reductions matched protein, with MSH3 and both wild-type and mutant HTT reduced ∼80-95% in striatum (Figure 1C-E, Supplemental Figure 2A,B) and medial cortex (Figure 1F-H, Supplemental Figure 2C,D), showing robust target knockdown across backgrounds (one-way ANOVA vs NTC, HTT p < 0.05, MSH3 p < 0.001).

**Figure 1:**
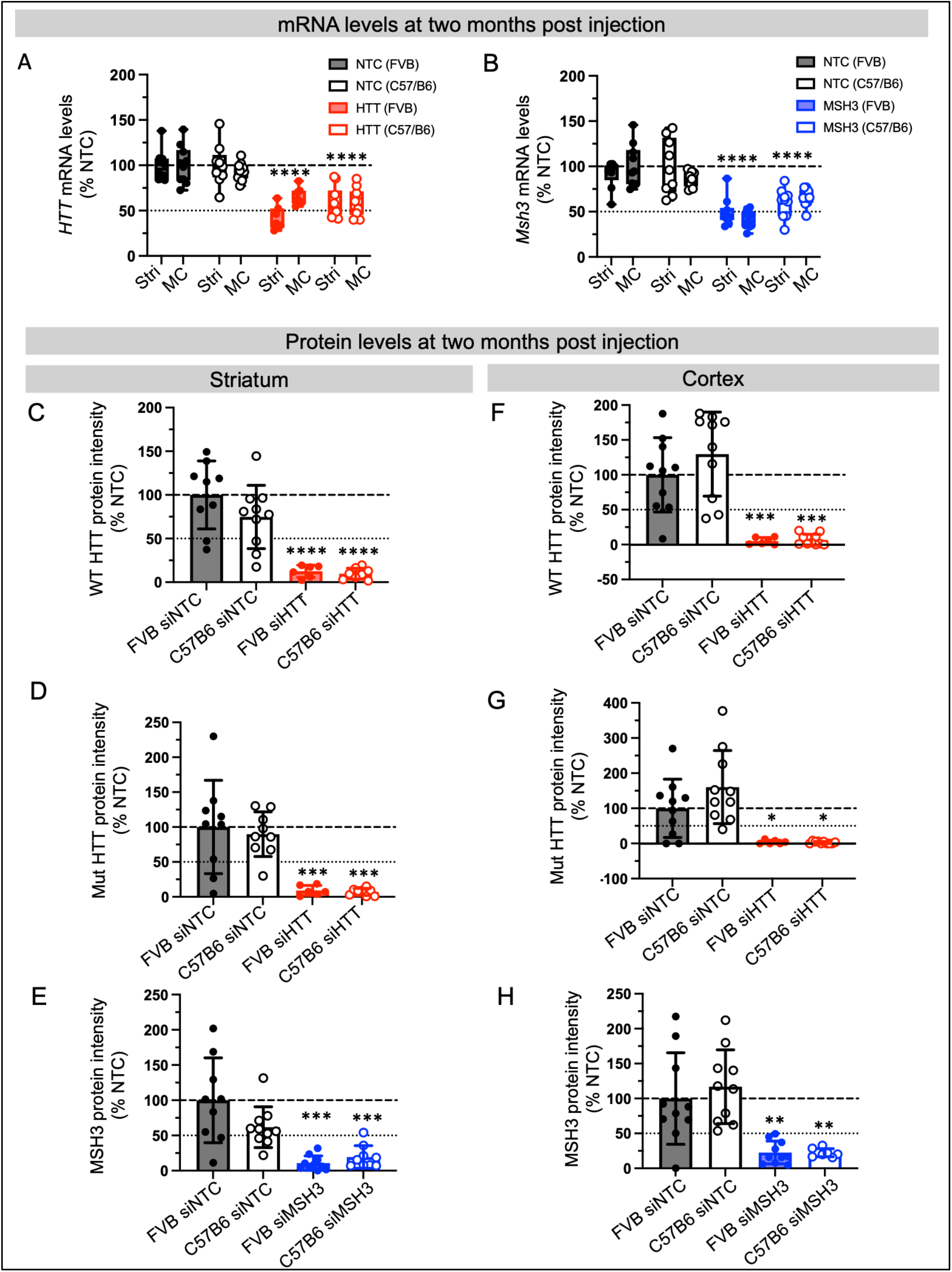
FVB and C57BL/6 BAC-CAG mice show robust HTT and *Msh3* mRNA and protein silencing after divalent siRNA ICV injection at 3 months, analyzed at 5 months. Solid circles: FVB; open circles: C57BL/6. (A) *Htt* mRNA (striatal left, medial cortex right). (B) *Msh3* mRNA. (C–E) Striatal protein: (C) wild-type HTT, (D) mutant HTT, (E) MSH3. (F–H) Medial cortex protein: (F) wild-type HTT, (G) mutant HTT, (H) MSH3. One-way ANOVA with Tukey post hoc vs FVB NTC. n = 7–10. *p<0.05, **p<0.01, ***p<0.001, ****p<0.0001.

**Figure 2:**
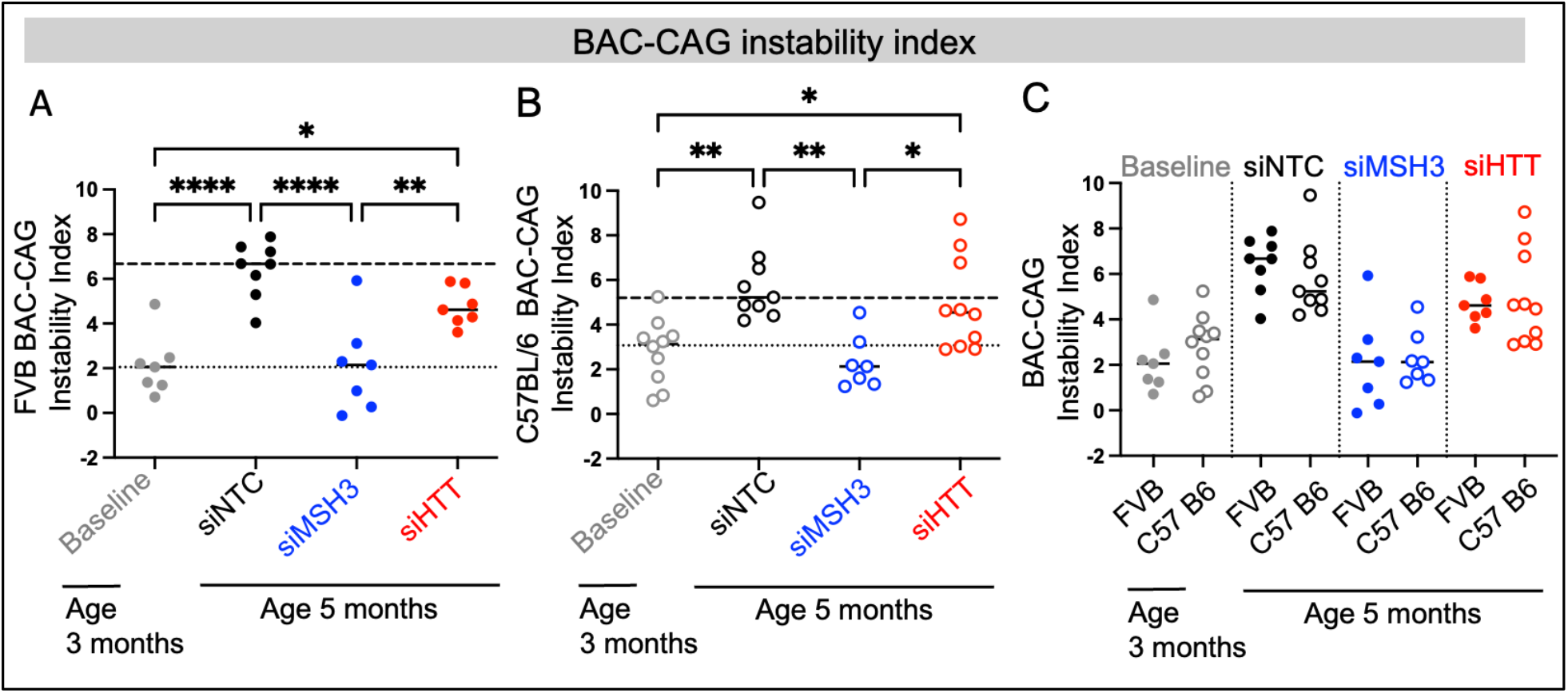
BAC-CAG mice on FVB and C57BL/6 backgrounds show robust somatic CAG expansion, blocked by MSH3 siRNA, unchanged by HTT lowering. (A) FVB somatic instability index. (B) C57BL/6 index. (C) FVB vs C57BL/6 comparison; short dashed line = median baseline (3 mo), long dashed = median NTC (5 mo). Solid: FVB, open: C57BL/6. Index calculated per Lee et al., 2010. n = 7–10. *p<0.05, **p<0.01, ***p<0.001, ****p<0.0001.

Somatic CAG repeat expansion in the striatum was measured using the somatic instability index, measuring expansion skew over time.^43^ A separate cohort was sacrificed at injection to define initial baseline instability (age three months). On average, the starting repeat size was similar between strains (Supplemental Figure 1, 131.4 for FVB and 130.1 for C57BL/6). At age five months, expansion indices were compared to baseline (age 3 months) and across treatments (NTC, MSH3-, and HTT-targeting siRNA). Both FVB and C57BL/6 BAC-CAG mice showed significant expansion over two months (Figure 2A,B, FVB baseline vs NTC p < 0.0001; C57BL/6 baseline vs NTC p < 0.01).

Notably, we observed that FVB mice showed lower baseline somatic instability at the time of injection (average baseline instability index = 2.1) and an increase of approximately five index units by five months of age (average NTC instability index at 5 months = 6.4). In contrast, C57BL/6 mice exhibited higher baseline instability (baseline instability index = 3.4) and a smaller increase over the same period, with an increase of approximately three instability index units (instability index at 5 months = 5.2). As a result, the overall delta of change in expansion over two months was reduced in the C57BL/6 background compared to FVB, despite statistically significant expansion in both lines. This reduced effect size in the C57BL/6 line may be important in future experiments when a power analysis for somatic expansion is performed.

Despite the difference in effect size, we observed that the somatic expression was blocked with MSH3-siRNA in both strains (Figure 2C, FVB siMSH3 instability index 2.08, one way NOVA vs NTC; C57/B6 siMSH3 instability index 2.3, one way ANOVA vs NTC, p < 0.001 both stains) and unchanged by HTT-siRNA in both strains (FVB siMSH3 instability index 4.8, one way ANOVA vs NTC; C57/B6 siMSH3 instability index 4.9, one way ANOVA vs NTC, p < 0.001 both stains) effects that have been previously observed in response to MSH3 or HTT modulation.^34, 35, 37, 44, 45^

Mismatch repair (MMR) genes have been shown to regulate the rate of somatic CAG repeat expansion in HD, and variation in expression of these factors can influence expansion dynamics.^4, 12, 29, 35, 37, 45^ To assess whether transcriptional differences, including MMR genes, explain strain-dependent effects, we performed RNA sequencing on striatal tissue from FVB and C57BL/6 BAC-CAG mice. Figure 3A shows a volcano plot of differentially expressed genes, most of which are pseudogenes, likely reflecting the use of the C57BL/6 genome as reference and strain-specific variation.^46^ Relative expression of *Htt* and MMR genes is shown in Figure 3B and Supplemental Figure 3, with pro- and anti-expansion genes in red and green, respectively. No significant differences were detected in endogenous *Htt*, human *HTT*, or MMR genes by RNA sequencing or Quantigene analysis (Figure 3B, Supplemental Figure 3). For MMR genes, *Msh2* was highest and *Pms1* was lowest expressed by mRNA levels in both strains (Figure 3B).

**Figure 3:**
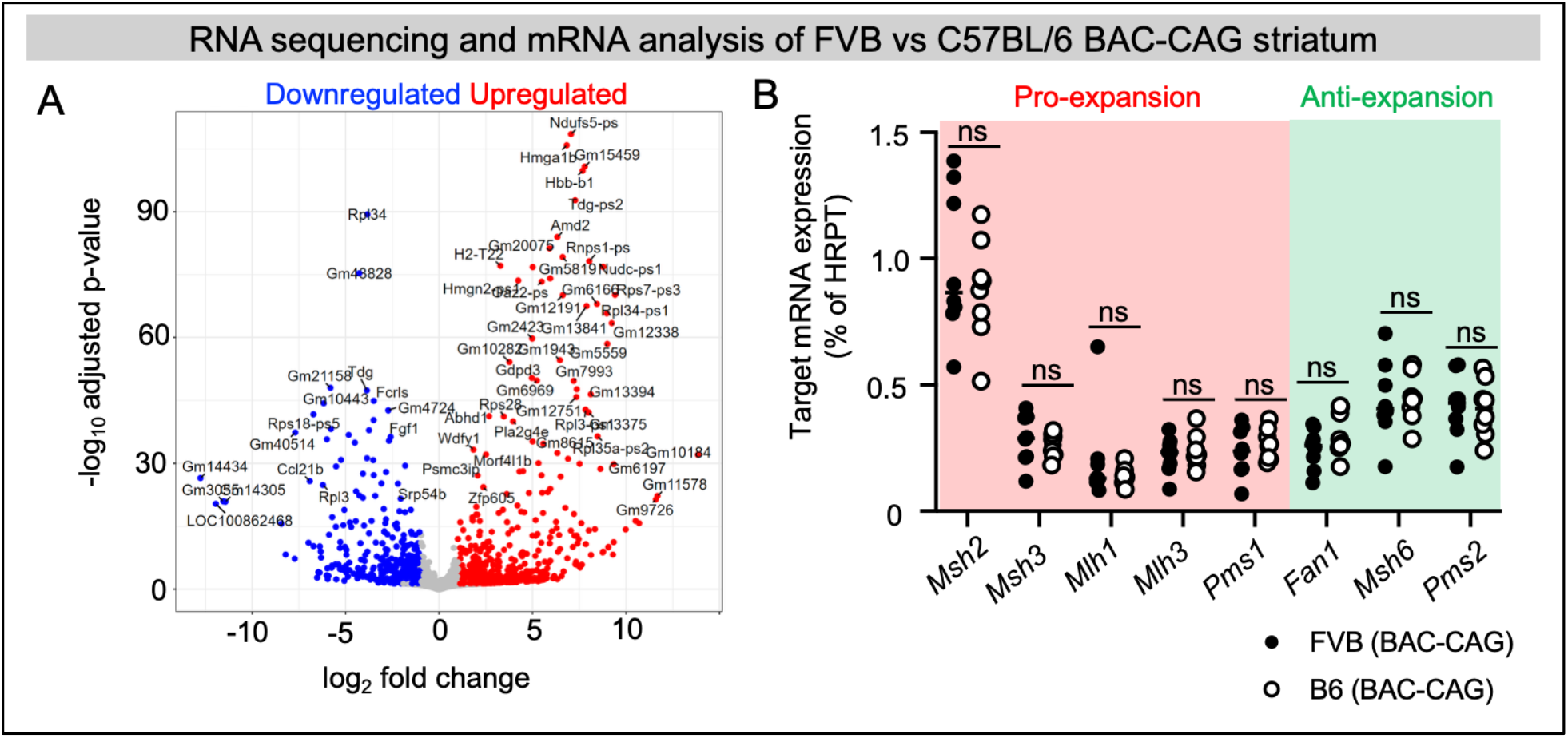
Strain-dependent RNA-seq differences in BAC-CAG striatum. (A) Volcano plot, red = up, blue = down in FVB vs C57BL/6. DESeq2-adjusted p-values shown (B) QuantiGene of MMR genes; solid = FVB, open = C57BL/6; genes categorized by effect on somatic instability. red = pro-expansion, green = anti-expansion; FVB vs C57BL/6. n = 7–10. *p<0.05.

These results show that both BAC-CAG strains respond to MSH3- and HTT-siRNA by target knockdown. Both strains show robust somatic CAG expansion, with FVB mice displaying slightly larger effects over two months, which may accumulate over time in longer studies. Further, both strains demonstrate blocked expansion to MSH3, but not HTT, silencing. Overall, both backgrounds provide reliable models for studying somatic expansion and therapeutic modulation in HD.

## DISCUSSION

In summary, we characterized somatic CAG repeat expansion in BAC-CAG mice on FVB and C57BL/6 backgrounds with and without siRNA-mediated modulation of HD modifiers, MSH3 and HTT demonstrating both models show robust, measurable somatic expansion over two months. mRNA expression of MMR and key HD associated genes were comparable between strains.

We did observe a non-significant increase in the magnitude of change in expansion over two months with an instability index change of +5 FVB versus +2 C57BL/6 which may impact sample size in further study design or accumulate over a multi-month long study. This observation suggests both strains of BAC-CAG mice can effectively model somatic expansion,^36^ though smaller effect sizes in the C57BL/G strain may require larger groups or longer observation periods (e.g., extending two-to three-month intervals) for adequate statistical power. Both strains responded robustly to MSH3 modulation, supporting their use in testing preclinical MSH3-targeted therapies (ASOs, siRNA, small molecules, AAV, etc.).^7, 18, 34, 35, 44, 45, 47, 48^ Though robust differences were not observed in this study, our work adds to prior literature demonstrating *Msh3* modulation has varying impacts on somatic expansion depending on the mouse model genetic background used.^10, 31^

Mouse models play a central role in preclinical HD research,^2, 5, 11, 27-29^ with varied models emphasizing distinct aspects of disease biology, including somatic expansion, HTT expression and isoforms, motor phenotypes, sleep disruption, and aggregate formation.^31, 33, 36^ Beyond somatic instability, HTT gene biology, including transcriptional expression, protein binding factors, and regulatory context, also shapes disease outcomes, independently or in combination with somatic expansion.^16, 49^ Human and mouse *HTT* genes differ in structure and regulation, including sequence, regulatory elements, protein binding sites, and intron-exon sequence and usage, which affect gene expression and processing.^50^ These human-mouse sequence differences can influence downstream HTT biology, further highlighting the need to carefully select and design models for preclinical studies.^49, 51^ For example, in humans and mouse models of HD, the presence and position of CAA interruptions at the end of the expanded CAG tract, along with intronic sequence context, for example influence somatic instability and disease phenotypes^4, 25, 52, 53^ The exact contribution of each of these *HTT* gene elements on HD progression in humans is not fully clear, making mouse models that can assess these features essential. Further, additional phenotypes not assessed in this study, such as aggregate formation, neuronal dysfunction or behavior, may also be important to consider in future studies using the BAC-CAG C57BL/6 strain.

Outside of strain background, other parameters may influence the comparison between the BAC-CAG and models such as the Q-series. For example, Q111 is a knock-in model in which the mouse *Htt* gene carries 111 CAG repeats within the endogenous locus on chromosome 4, maintained on a C57BL/6 background.^54^ As a knock-in model, Q111 expresses an expanded version of the mouse *Htt* in exon 1 within the native mouse *Htt* regulatory context.^54^ These design differences (knock-in vs transgenic) can influence gene expression, regulatory interactions, phenotype, and mRNA and protein dynamics when modeling human disease.^55, 56^

Limitations of this work include that while bulk tissue analyses are informative, dissecting selective neuronal vulnerability due to somatic expansion in HD will likely require single-cell approaches, which is ongoing.^6, 20^ We also characterized mRNA expression of MMR genes. However, mRNA levels do not always match protein levels, especially in DNA repair pathways.^57-59^ Protein abundance and the relative ratios protein expression within the MMR system may better reflect the contributions of MMR pathway to disease. Proteomic profiling will therefore be important to fully define these effects.

These data provide a foundation for comparing C57BL/6 BAC-CAG mice with knock-in Q-length models such as Q111 and Q140. Characterizing the BAC-CAG line, which carries the full human expanded HTT gene within the human HTT regulatory context, on the same strain as the Q-series, now allows these questions to be explored. Overall, this work advances preclinical studies in somatic expansion-targeted therapeutics and supports the translation of findings from mouse models to individuals living with HD.

## Author Contributions

Conceptualization: AK, NA, DH, TFV, JB; Methodology: AK, NA, MF, DH, TFV, XWY; Investigation: JB, AS, SH, ES, EL, NY, DO; Formal analysis: JB, ES, SH, AK, Writing – Original Draft: JB, AK; Writing - Review & Editing: JB, AS, SH, ES, EL, NY, DO, DH, MF, NA, AK, XWY; Visualization: JB, AK, ES, SH; Resources: DH, MD, NA, AK, XWY; Supervision: DH, MD, NA, AK; Funding acquisition: AK, NA, MD

## Funding Acknowledgements

The authors thank the CHDI foundation for the JSC A-5038 and A-17281 awards to AK, NA, MD. The authors thank the National Institutes of Health for awards S10OD020012 and S10OD036329 to AK, and F31 NS132424 to JNB.

## Data availability

Study data are available upon request to the corresponding author. Materials and resources used in this study are publicly or commercially available.

