## Supplemental Material for "C57BL/6 BAC-CAG Huntington’s disease mice show somatic CAG expansion and responses to small interfering RNAs comparable to the FVB strain"

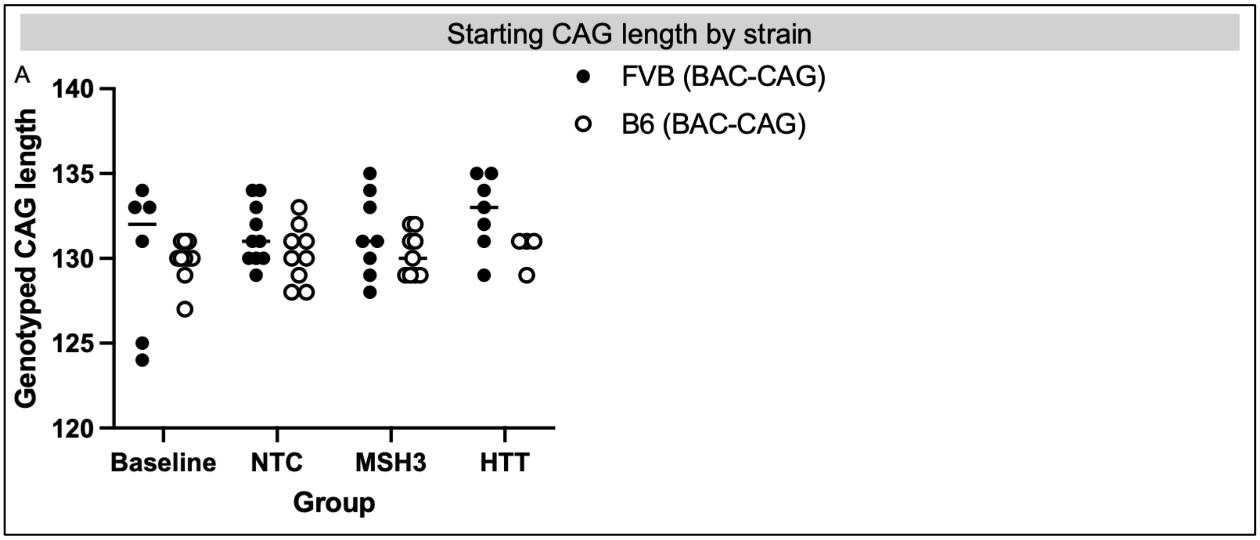


**Supplemental Figure 1:** Inherited CAG repeat lengths were balanced across treatment groups. Genotypes and CAG levels were determined from tail samples.

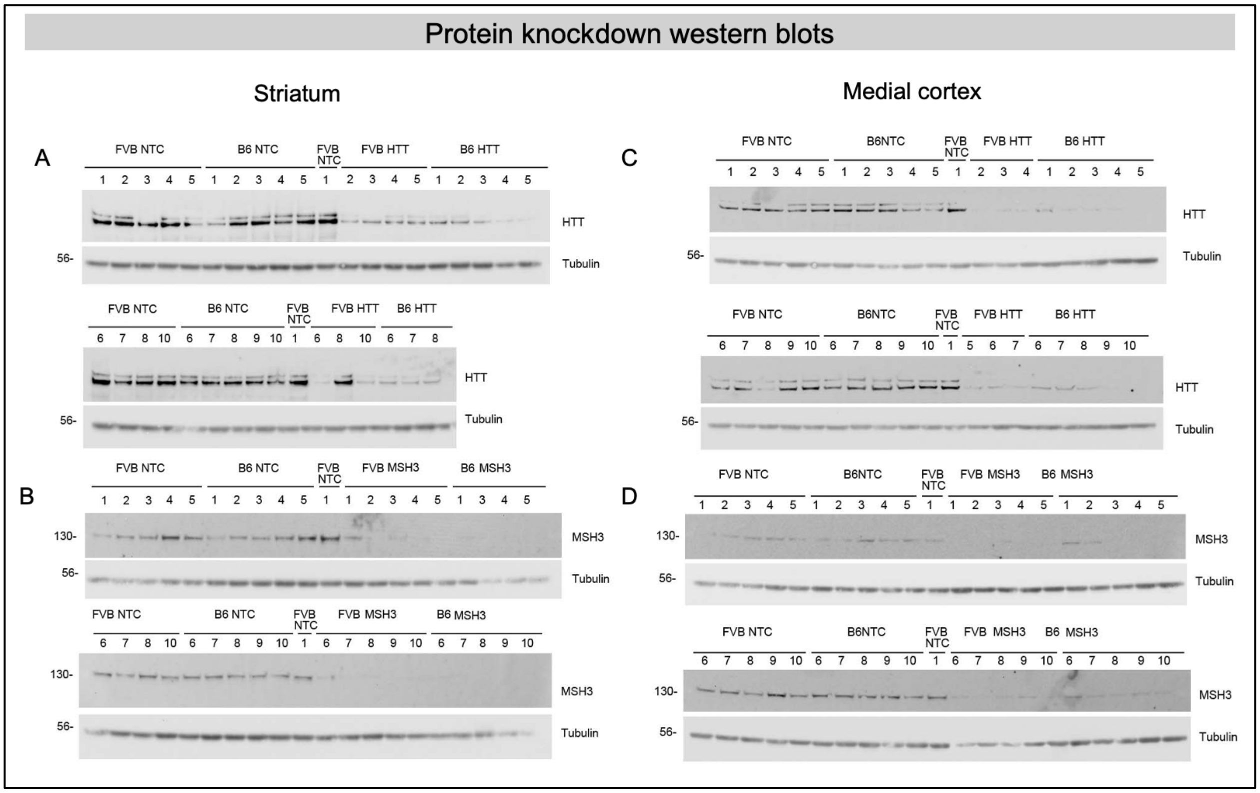


**Supplemental Figure 2:** Western blots used to quantify protein levels of HTT and MSH3 at two-months post-siRNA injection. (A-B) Striatum levels of (A) HTT and mutant HTT, or (B) MSH3 in FVB and C57BL/6 mice. (C-D) Medial cortex levels of (C) HTT and mutant HTT, or (D) MSH3 in FVB and C57BL/6 mice. Mice were treated with non-targeting control (NTC), *Msh3*-targeting, or *HTT*-targeting siRNA. n= 7-10 per group.

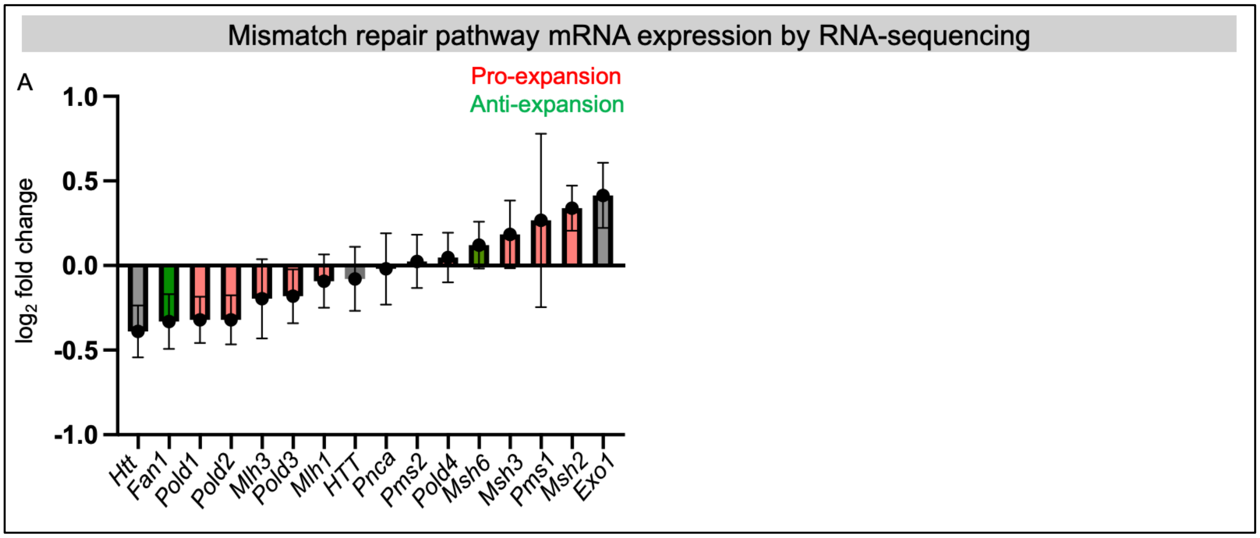


**Supplemental Figure 3:** HD modifier and mismatch repair pathway expression by RNA-sequencing (B) Relative expression of selected HD genes; red = pro-expansion, green = anti-expansion; FVB vs C57BL/6. DESeq2-adjusted p-values shown. (C) QuantiGene of MMR genes; solid = FVB, open = C57BL/6; genes categorized by effect on somatic instability. n = 7–10. *p<0.05.

**Supplemental Table 1:** siRNA sequences used in vivo

**Supplemental Table 2:** RNA-sequencing comparison between the striatum's of FVB and C57BL/6 strains of BAC-CAG mice.
